# Extreme stiffness of neuronal synapses and implications for synaptic adhesion and plasticity

**DOI:** 10.1101/2020.08.31.273466

**Authors:** Ju Yang, Nicola Mandriota, Steven Glenn Harrellson, John Anthony Jones-Molina, Rafael Yuste, Roger Lefort, Ozgur Sahin

## Abstract

Synapses play a critical role in neural circuits, and they are potential sites for learning and memory. Maintenance of synaptic adhesion is critical for neural circuit function, however, biophysical mechanisms that help maintain synaptic adhesion are not clear. Studies with various cell types demonstrated the important role of stiffness in cellular adhesions. Although synaptic stiffness could also play a role in synaptic adhesion, stiffnesses of synapses are difficult to characterize due to their small size and challenges in verifying synapse identity and function. To address these challenges, we have developed an experimental platform that combines atomic force microscopy, fluorescence microscopy, and transmission electron microscopy. Here, using this platform, we report that functional, mature, excitatory synapses had an average elastic modulus of approximately 200 kPa, two orders of magnitude larger than that of the brain tissue, suggesting stiffness might have a role in synapse function. Similar to various functional and anatomical features of neural circuits, synaptic stiffness had a lognormal-like distribution, hinting a possible regulation of stiffness by processes involved in neural circuit function. In further support of this possibility, we observed that synaptic stiffness was correlated with spine size, a quantity known to correlate with synaptic strength. Using established stages of the long-term potentiation timeline and theoretical models of adhesion cluster dynamics, we developed a biophysical model of the synapse that not only explains extreme stiffness of synapses, their statistical distribution, and correlation with spine size, but also offers an explanation to how early biomolecular and structural changes during functional potentiation could lead to strengthening of synaptic adhesion. According to this model, synaptic stiffness serves as an indispensable physical messenger, feeding information back to synaptic adhesion molecules to facilitate maintenance of synaptic adhesion.

---

Synapses are highly-specialized intercellular junctions comprised of pre- and postsynaptic structures that are tightly connected by synaptic adhesion molecules^1^. Because synapses play a critical role in learning and memory, the biochemical, morphological and electrophysiological characteristics of synapses have been widely investigated. Due to their role in adhesion of pre- and postsynaptic cells, mechanics of synapses can also be important for synapse function^2^. Indeed, studies with various cell types have demonstrated the important role played by the mechanical stiffness of adhering structures on the strength of cellular adhesion^3-6^. It could similarly be the case that stiffness of synapses affects the strength of adhesion between pre-and postsynaptic structures. Understanding factors that control synaptic adhesion is important due to its role in maintaining connectivity and function of neural circuits. However, studying mechanical characteristics of synapses are made difficult by the small sizes and complex sub-structures of synapses, as well as the need for accurately identifying synapses, which require biochemical and ultrastructural analysis alongside mechanical measurements.

We have developed an experimental platform to carry out mechanical, biochemical and ultrastructural analysis of synapses by combining atomic force microscopy (AFM), fluorescence microscopy and transmission electron microscopy (TEM). Although AFM has been previously used to probe mechanics of synapses^7^, we used a recently developed cell stiffness imaging method^8^ based on torsional harmonic AFM (TH-AFM)^9,10^ which offers high-resolution, high-speed, quantitative stiffness mapping of live cells^8^, making the method suitable for quantifying stiffness mapping of a large number of synapses. We combined this AFM method with fluorescence microscopy to identify mature synapses and monitor their activity, and with TEM to reliably identity synapses by observing the synaptic cleft, presynaptic vesicles, and postsynaptic density, and also to correlate AFM stiffness images to synaptic ultrastructure. Although AFM is commonly used in combination with fluorescence microscopy to study biological cells, combined use of AFM and TEM has been limited to imaging of filamentous biomolecules deposited on substrates^11,12^. To achieve combined AFM and TEM imaging on the same individual synapse, AFM stiffness mapping of the delicate synapses has to be performed without compromising their structural integrity prior to TEM analysis. The TH-AFM allows gentle probing of synapse stiffness with low peak forces during the tapping-mode AFM imaging. To facilitate combined TH-AFM and TEM imaging, we developed protocols for preparing TH-AFM-imaged samples for serial-sectioning, TEM imaging, and for alignment of AFM and TEM data (See Methods).

With our experimental setup, we measured the elastic modulus of hundreds of live mature excitatory synapses and mapped stiffness of postsynaptic spines and presynaptic boutons identified by TEM. These experiments showed that spines are two orders of magnitude stiffer than the average brain tissue. Localization of such high stiffness to a submicron structure could indicate that stiffness of spines might have an important role in synaptic function. Additional observations of the statistical distribution of elastic modulus values, correlation between stiffness and spine size, and a mechanical model of highly crosslinked actin network provided further clues to the physical origin and biochemical regulation of spine stiffness, and suggested that the spine stiffness could be a quantity that adapts to the synaptic function. Based on stiffness-dependent stochastic dynamics of adhesion clusters^13,14^, we developed a simple biophysical model of the synapse that points to an indispensable role of stiffness in synapse function. According to this model, functional potentiation of the synapse leads to increased stiffness, which, in turn, results in a stronger synaptic adhesion.

## Results

### Live nanomechanical imaging of cultured neurons

We characterized the nanomechanical properties of live hippocampal neurons on 14-25 days *in vitro* (DIV) using TH-AFM (Fig. 1a). TH-AFM uses a T-shaped cantilever with an offset tip (Fig. 1b) and detects both vertical and torsional deflection signals of the cantilever during scanning of the sample. The vertical signal is used for the height feedback to generate the topographical image and the torsional signal provides instantaneous force to calculate mechanical properties during the imaging process^9^ (Fig. 1c). To minimize forces acting on delicate neural structures, we used low peak tapping forces (around 300 pN).

**Figure 1:**
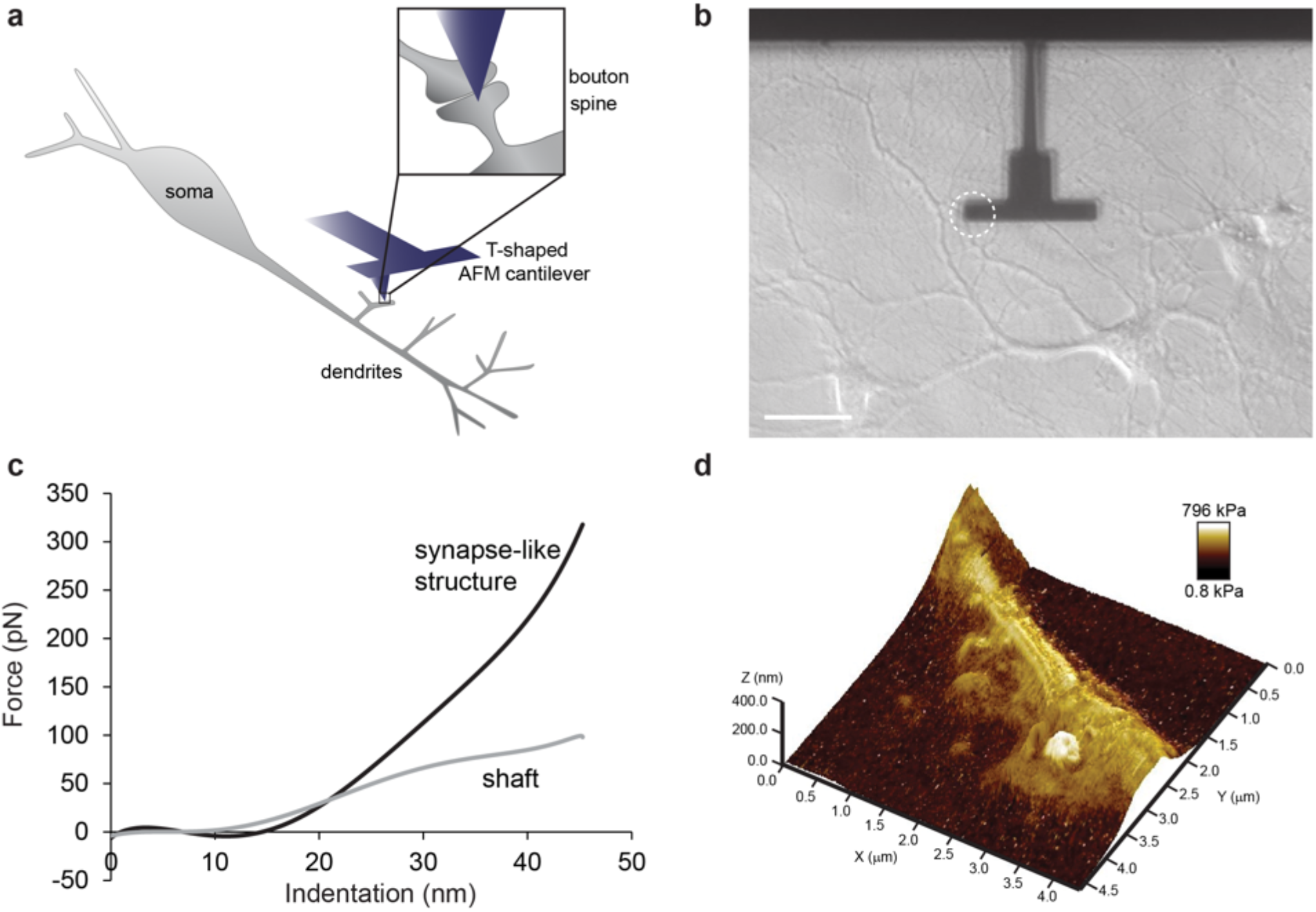
Nanomechanical imaging of live hippocampal neurons with TH-AFM. **a** A schematic diagram illustrating the set-up of the nanomechanical imaging platform. Neurons were cultured in a glass bottom dish. TH-AFM imaging was performed by scanning a sharp tip across the sample surface while the interaction force between the tip and the sample was monitored. The zoomed-in inset illustrates the AFM tip over a synapse. **b** Optical image of a T-shaped cantilever over a neuron culture. The tip was facing towards the culture dish, and was placed on the left side of the cantilever (indicated by the dashed circle). Scale bar: 40 μm. **c** Representative force-distance curves of a synapse-like structure (black) and a dendritic shaft (grey). See Supplementary Fig. 1 for the corresponding AFM images. **d** Three-dimensional topographical rendering of a synapse-like structure with the color indicating elastic modulus in log scale. The elastic modulus values of the stiff structure and the shaft are 626.8 kPa and 38.6 kPa, respectively.

We cultured rat hippocampal neurons in glass bottom dishes and imaged live neurons in Tyrode’s buffer at room temperature. To capture the corresponding optical images, we placed the TH-AFM apparatus above an inverted optical microscope with high magnification and high numerical aperture objectives (see Methods). Hippocampal neurons were characterized as large pyramidal cell bodies with long axons and branched dendritic shafts, forming complex neurite networks (Fig. 1b). We optimized the culture density so that individual synapses could be physically accessed by the TH-AFM tip.

Our imaging technique generated topographical and mechanical images simultaneously during live imaging at nanometer resolution. A three-dimensional topographical rendering of a synapse-like structure with the color indicating elastic modulus is shown in Fig. 1d. Within the network of processes, we observed surprisingly stiff synapse-like round structures near compliant neurites. We used the following criteria for visually-identified synapse-like structures under TH-AFM: (i) distinct structures with stiffness over 20 kPa in the mechanical image, (ii) height below 1.5 μm and (iii) in close proximity (within 2 μm) to a nearby neurite in the topographical image. Based on these criteria, we acquired images of hundreds of synapse-like structures and nearby shafts (see Methods for details of quantitative stiffness measurements), and found 77.8% of synapse-like structures had a stiffness over 100 kPa. These stiff structures usually displayed a round shape in topographical images and a dark contrast in optical images (Fig. 2a). Based on their morphology and proximity to neurites, we hypothesized that these stiff structures could be synapses.

**Figure 2:**
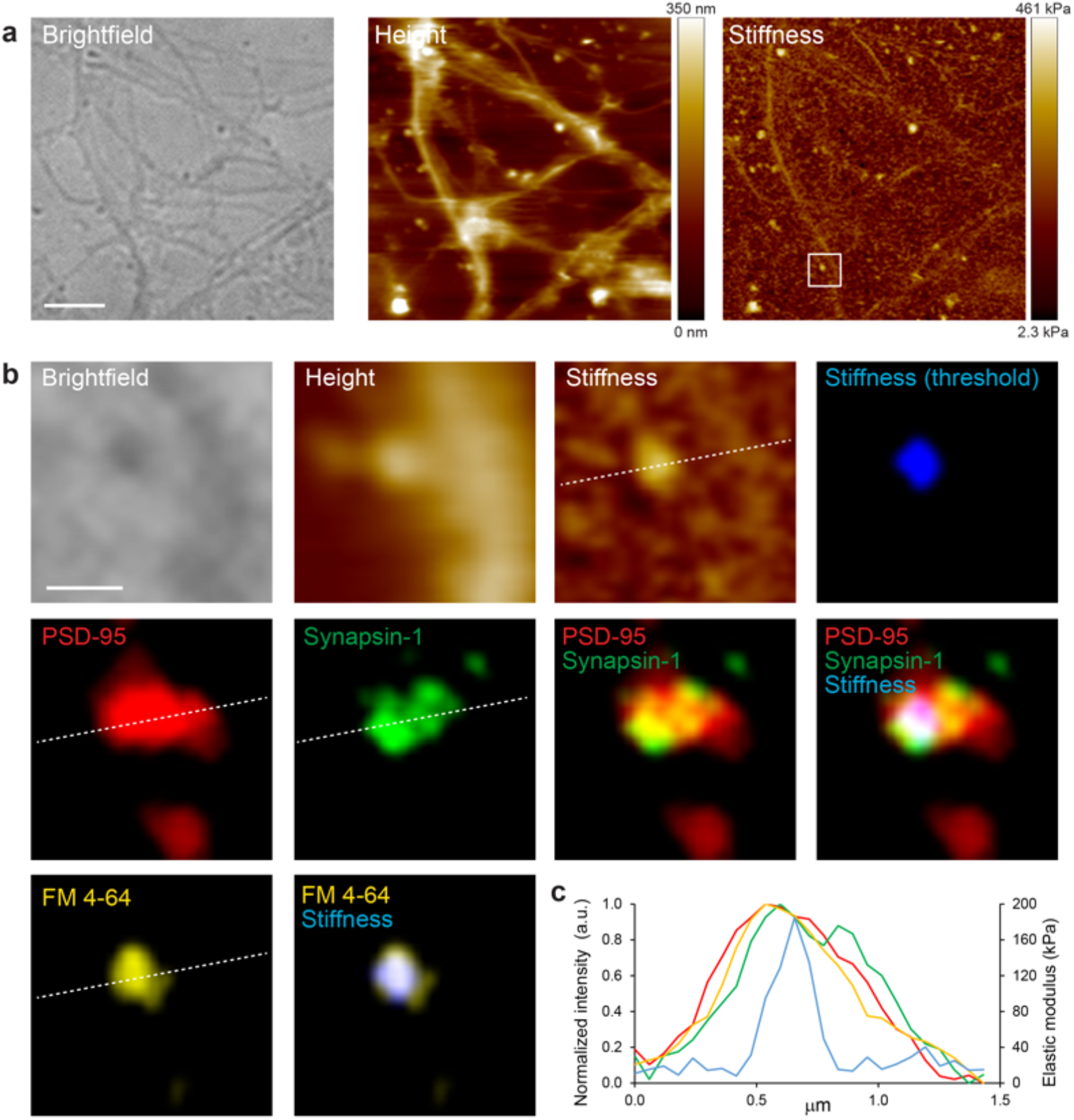
Correlative TH-AFM/fluorescence imaging shows stiff synapse-like structures are functional mature excitatory synapses. **a** Aligned brightfield, AFM height, and AFM stiffness images of the same area in live neuron cultures. The color in AFM images represents height (linear scale) and elastic modulus (log scale), respectively. Scale bar: 3 μm. **b** Aligned brightfield, AFM height, AFM stiffness and fluorescence images of a representative stiff synapse from the boxed area shown in the stiffness image in **a**. After TH-AFM imaging, neurons were fixed and stained with postsynaptic marker PSD-95 (red) and presynaptic marker Synapsin-1 (green). Colocalization of these two markers identified mature synapses. Threshold was applied to the stiffness image colored in blue. An overlay image of stiffness, PSD-95 and Synapsin-1 shows the stiff structure was a mature synapse. In addition, neurons were stained with FM 4-64 (yellow) after TH-AFM imaging to label functional synapses. Scale bar: 500 nm. **c** Fluorescence intensity and elastic modulus profiles along the dashed lines over the synapse in **b** show that high stiffness (blue) overlaps with synaptic markers, PSD-95 (red), Synapsin-1 (green), and FM 4-64 (yellow). Note that the elastic modulus has a narrower peak than fluorescence signals.

### Stiff synapse-like structures are functional mature excitatory synapses

To verify the identity of stiff structures revealed by TH-AFM, we performed immunostaining against various synaptic markers after TH-AFM imaging and correlated immunofluorescence images with AFM images. Figure 2b shows a representative stiffness image and the corresponding immunostaining results of a synapse. We defined mature excitatory synapses as the colocalization of the presynaptic marker Synapsin-1, and the postsynaptic marker PSD-95 (see Methods for colocalization detection). We found that all stiff spherical structures revealed by TH-AFM were co-labeled with both Synapsin-1 and PSD-95 (Supplementary Fig. 2a, b), suggesting that they were mature excitatory synapses. Interestingly, time-lapse TH-AFM imaging of the same synapse showed that synapse stiffness did not vary significantly during imaging (Supplementary Fig. 3).

To further characterize these synapses and monitor their activity, we incubated live neurons with FM 4-64 dye after TH-AFM imaging (Fig. 2b, Supplementary Fig. 2c, d). FM dye binds to cellular membrane and gets internalized after neurotransmitter is released, and thus only labels functional presynaptic boutons^15^. The results show that stiff structures were colocalized with FM puncta as well, indicating that these synapses were functional. Fluorescence intensity and elastic modulus profiles (Fig. 2c) show that high stiffness overlaps with synaptic markers and FM dye and that elastic modulus has a narrower peak than fluorescence signals. Taken together, the data confirm that stiff structures identified in cultured neurons were functional and mature excitatory synapses.

### Pre- and postsynapses have different stiffness

To distinguish between pre- and postsynapses at high resolution and to investigate whether they are mechanically distinct, we performed TEM imaging after TH-AFM. To locate the same synapses under TEM, we cultured neurons in homemade glass bottom dishes with photoetched gridded coverslips and used the numeric pattern as landmarks to align TEM images with optical and AFM images^16^ (Supplementary Fig. 4). Serial section TEM can reveal the ultrastructure of synapses at nanometer resolution^17^. We found that stiff structures were featured with presynaptic vesicles and postsynaptic density (Fig. 3 and Supplementary Fig. 5). In particular, spines, rather than boutons, overlapped with high stiffness pixels. We conclude that the high stiffness observed under TH-AFM originated from postsynaptic spines and that postsynaptic spines are mechanically different from presynaptic boutons.

**Figure 3:**
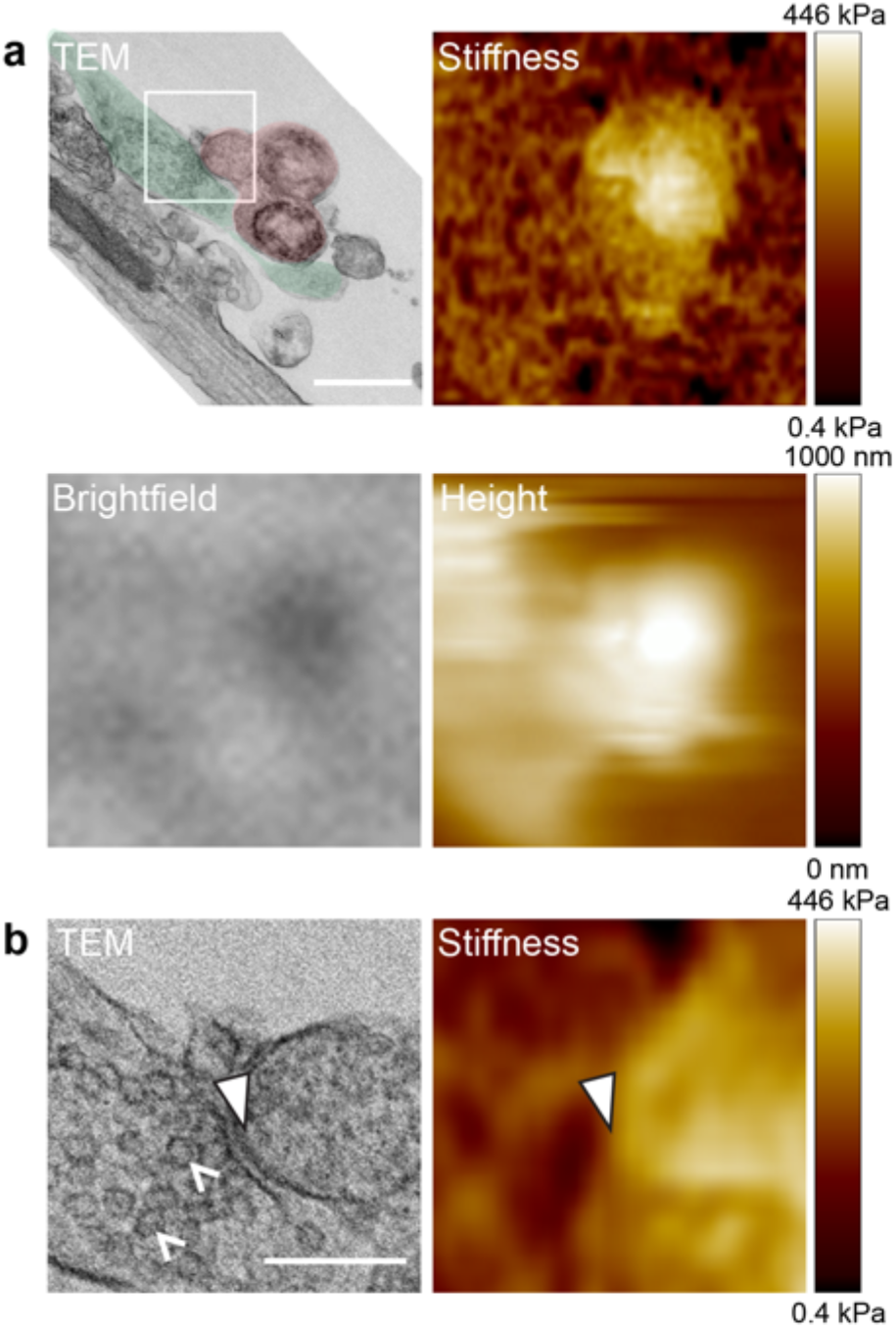
Correlative TH-AFM/TEM reveals ultrastructure of stiff synapses. **a** Aligned TEM, AFM stiffness, brightfield, and AFM height images of the same synapse. We used the numeric pattern on photoetched coverslips as landmarks to align TEM images with optical and AFM images. In the TEM image, the bouton is shaded in green and the spine is shaded in red. High stiffness in the AFM stiffness image overlapped mostly with the spine head (209 kPa), while the bouton and dendritic shaft showed lower stiffness. The stiff spine displayed distinct contrast in the optical brightfield image and topographical features in the AFM height image. Scale bar: 500 nm. **b** Zoomed-in TEM and AFM stiffness images of the boxed area shown in the TEM image in **a** illustrates the synaptic cleft. White arrowheads point to the postsynaptic density. White carets point to presynaptic vesicles. Scale bar: 200 nm.

### Spines are substantially stiffer than shafts

Intriguingly, the stiffness of spines fell within a wide range. The stiffness of shafts, on the contrary, was constantly low (Fig. 4a). Spines were on average 10 times stiffer than nearby shafts. The minimum, maximum, median, and mean elastic modulus values of spines are 23.2 kPa, 671.9 kPa, 166.9 kPa and 201.3 kPa, whereas the minimum, maximum, median, and mean elastic modulus values of shafts are 7.1 kPa, 67.4 kPa, 20.7 kPa and 23.6 kPa (see Methods). The distributions of spine and shaft stiffness were strongly skewed with heavy tails and exhibited the characteristics of a lognormal distribution. We measured the apparent spine size from AFM stiffness images by thresholding the stiffness signal to identify region of interest for area measurement (see Methods). We found that the distribution of apparent spine size was also skewed with a heavy tail (Fig. 4b). Spine stiffness measurements were positively correlated with apparent spine size (Fig. 4c) (see Supplementary Notes for additional discussion about the sensitivity of the correlation analysis to the contact mechanics models), suggesting that larger spines may have underlying changes in actin architecture^18^. In addition, we found that immature protrusions, likely filopodia, were not stiff (Supplementary Fig. 6), suggesting that stiffness may increase as spines mature, and possibly change with synaptic function and activity. The surprisingly high stiffness of spines may represent a unique parameter related to synaptic activity and function that is complementary to the well-established biochemical and electrophysiological parameters.

**Figure 4:**
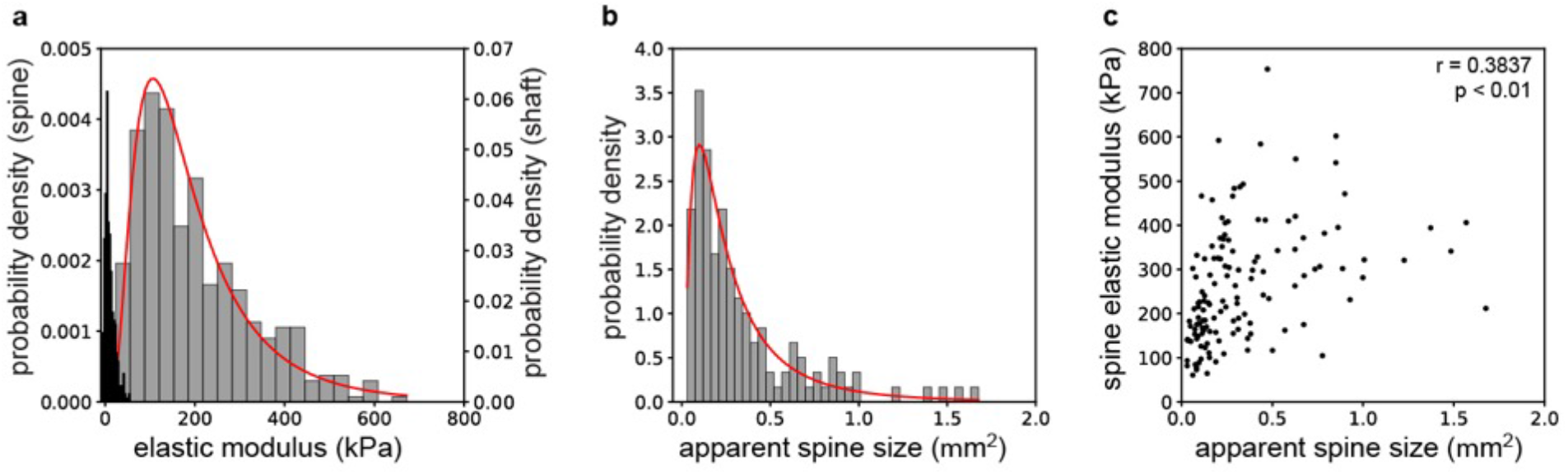
Spine stiffness is higher than shaft stiffness, and is correlated with spine size. **a** Histogram of elastic modulus of spines (grey) and nearby shafts (black) with bins = 20. n = 409 spines / 30 neuron cultures. We fitted the logarithm data with a normal distribution (spine stiffness fitted curve shown here in red). The mean elastic modulus values of spines and shafts from the fitting are 164.3 kPa and 21.4 kPa, respectively. We tested the goodness of fit using Kolmogorov-Smirnov tests (p_spine_ = 0.3192, p_shaft_ = 0.2446). **b** Histogram of the apparent spine size measured from AFM stiffness images with bins = 37. n = 134 spines / 20 neuron cultures. We fitted the logarithm of data with a normal distribution (red fitted curve with a mean spine size of 0.23 μm^2^.) and tested the goodness of fit using a Kolmogorov-Smirnov test (p = 0.9486). **c** Correlation of spine elastic modulus with apparent spine size from **b**. r: Pearson correlation coefficient. p: significance of correlation with a two-tailed t test. p = 4.719E-06.

Our criteria for synapse-like structures under TH-AFM might exclude spiny synapses with a stiffness below 20 kPa or without distinct height features in topographical images (for example, spines buried underneath neurites), and shaft synapses that are formed directly on the shaft without protrusive spine structures. Nevertheless, given that the measured spine stiffness had a lognormal-like distribution with a peak around 164 kPa, very few spines would be excluded by our criteria.

### Mechanical synaptic plasticity

The average value of elastic modulus of spines (201.3 kPa) is approximately two orders of magnitude larger than the average elastic modulus values reported for the gray and white matter brain tissue^19^. The substantially high stiffness of spines suggests a potential novel physiological role. Stiffness could be necessary to maintain spine morphology in the presence of synaptic adhesion. The unique spine morphology with an enlarged head and a thin neck enables compartmentalization of chemical and electrical signals, which is critical for synaptic function and plasticity^20^. Treating the synapse as two soft bodies in adhesive contact suggests that spine stiffness has to be on the order of 100 kPa to maintain its shape (see modeling and calculations in Supplementary Notes and Supplementary Table 1 - 4), which is consistent with the measured values of spine stiffness in Fig. 4a. However, the stiffness requirement for maintaining spine morphology only specifies an approximate minimum value for stiffness, therefore it would not lead to a particular distribution of stiffness values, nor would it predict a positive correlation with spine size, which are clearly observed in our measurements in Fig. 4.

A potentially critical role for stiffness in synapse function could be due to the connection between stiffness and cell adhesion, which could explain both our quantitative measurements and the observed distributions and correlations. Previous studies of the stochastic dynamics of adhesion molecules^13^ clustered at the site of adhesion has shown that the overall lifetime of a cluster depends on the stiffness of adhering surfaces^14^. In this view, due to the stochastic nature of molecular interactions, adhesion bonds rupture and rebind continuously. When bonds are ruptured, the surfaces can deform due to a small but non-zero force that pulls the surfaces apart. As illustrated in Fig. 5a, rupture of bound molecules will increase the distance between those molecules, reducing the probability of their future rebinding, and thereby decreasing the lifetime of the adhesion cluster. When the surfaces get stiffer, the probability of rebinding increases, leading to stabilization of the adhesion cluster. Gao et al.^14^ showed that a typical adhesion cluster is substantially stabilized as the sample stiffness increases beyond 50 to 100 kPa (Fig. 5b). The stiffness-dependent lifetime of adhesion clusters could also be applicable to synaptic adhesion, because pre- and postsynapses are connected together by transmembrane synaptic adhesion molecules^1^. Indeed, one of these molecules, N-cadherin, has been shown to be strengthened on stiffer substrates in C2 mouse myogenic cells^6^. Importantly, our measured spine elastic modulus values in Fig. 4a correspond to the regime where the lifetime of adhesion would be greatly enhanced, as shown in Fig. 5b.

**Figure 5:**
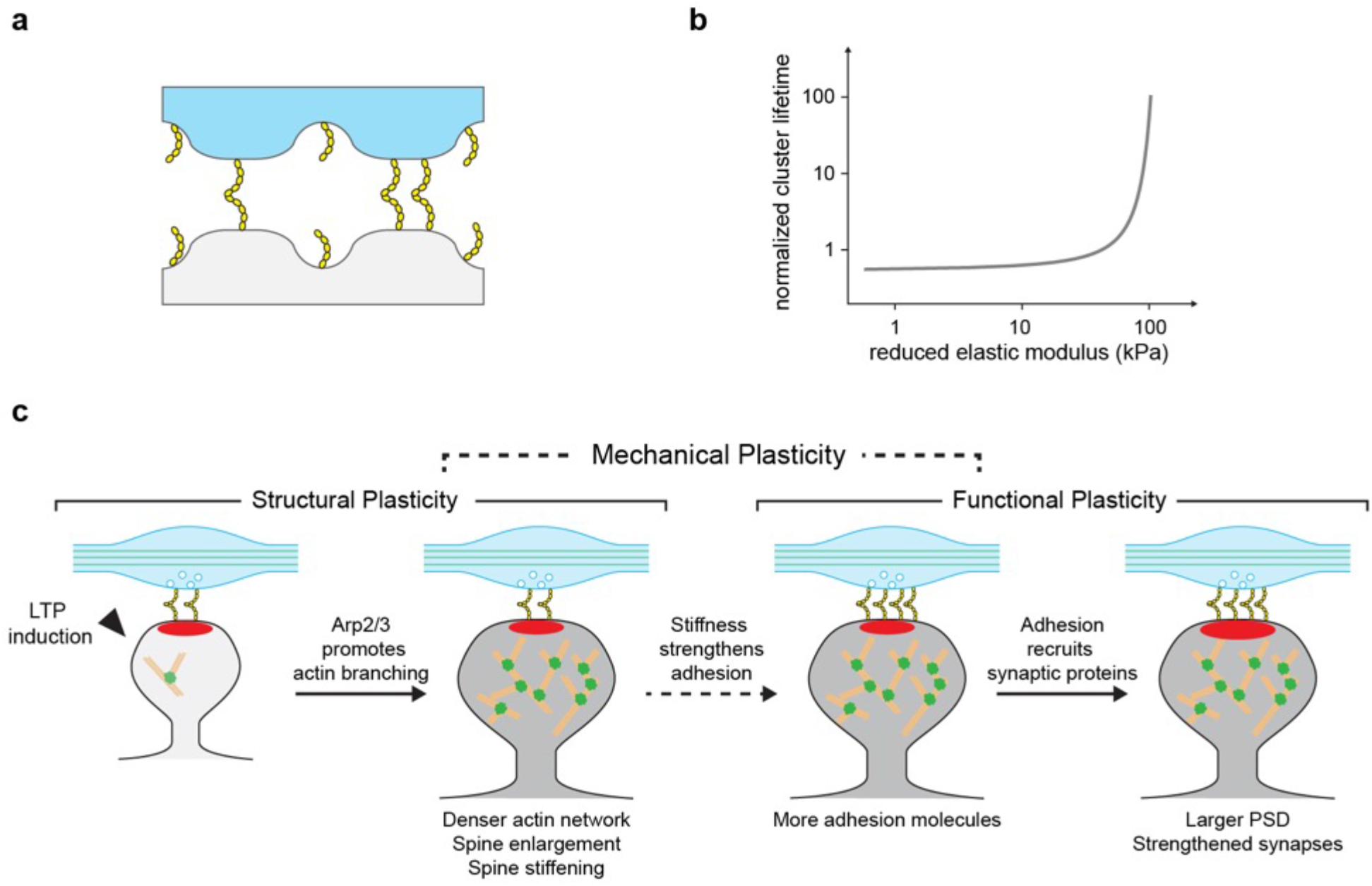
Mechanical synaptic plasticity. **a** Two elastic bodies (blue and grey) are connected by a cluster of adhesion molecules (depicted in yellow). Following the rupture of bonds between individual molecules, elastic recoil reduces the probability of rebinding due to the separation of binding molecules, causing stochastic dynamics of the adhesion cluster. **b** The lifetime of a cluster increases drastically as the sample stiffens. Diagrams in (**a**,**b**) are adapted from Qian and Gao^14^. **c** The mechanical synaptic plasticity model based on the stiffness-dependent stochastic dynamics of adhesion clusters. The schematic illustrates a simplified sequence of synaptic changes during LTP. The postsynaptic spine (grey) and the presynaptic axonal bouton (blue) are physically connected by synaptic adhesion molecules such as N-cadherin (yellow). Spine with postsynaptic density (PSD; red) has enriched actin networks (orange) cross-linked by Arp2/3 (green), while axon contains rigid microtubules (green lines). During LTP, spine exhibits both structural plasticity and functional plasticity with spine enlargement, actin reorganization, PSD enlargement, and increase of synaptic transmission. As shown by the dashed arrow, increased stiffness during structural plasticity (indicated by the darkening grey color of the spine) could enhance the adhesion at the synapse by increasing the surface density of bound adhesion molecules at the synapse. Adhesion molecules then help recruit synaptic proteins including PSD-95, AMPAR, and NMDAR, resulting in PSD enlargement and functional potentiation. Upon subsequent stimulations, a strengthened spine would have more Ca^2+^ influx, causing further spine stiffening and thus creating a positive feedback to maintain spine stiffness and synaptic potentiation.

Given the connection between stiffness and adhesion strength, we propose a biophysical model, the mechanical plasticity model, to explain how stiffness can be enhanced during long-term potentiation (LTP), and how the enhanced stiffness can facilitate maintenance of synaptic adhesion (Figure 5c). This role of spine stiffness fits well into the previously established LTP timeline: During LTP, NMDAR-dependent Ca^2+^ signaling cascade leads to structural plasticity involving spine enlargement and actin remodeling^21,22^, followed by an increase in adhesion molecules such as N-cadherin^23^, followed by enlargement of the postsynaptic density (PSD)^18,24^, and later an increase in synaptic receptors and enhanced synaptic strength. The structural changes that occur early in the LTP timeline, particularly the actin remodeling can lead to a substantial increase in the stiffness of actin network, which, in turn, would strengthen the synaptic adhesion.

High resolution imaging methods show that spines contain densely crosslinked actin networks^25,26^ (see also Supplementary Fig. 7), which could explain the origin of high stiffness of spines. To test this possibility quantitatively, we used a simple mechanical model of a three-dimensional network of fibers that relates the elastic modulus of the network to the geometry of the network with the following relationship^27^:

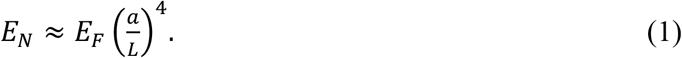

Here *E_N_* is the elastic modulus of fiber network, *E_F_* is the elastic modulus of individual fibers, *a* is fiber radius, and *L* is the distance between nodes of the network where fibers are connected. In the case of a crosslinked actin network, *a* would correspond to the nominal radius of actin filaments, and *L* would correspond to the average distance between crosslinks. Using and *a* ≈ 3.5 nm and *E_F_* ≈ 1GPa for the nominal radius and elastic modulus^28^ of F-actin filaments, Eq. (1) predicts that the average distance between crosslinks of the actin network of the spine to be in the range of 22 to 53 nm to match the experimentally observed elastic modulus values (i.e. 20 to 600 KPa). These number agree with electron microscopy images of actin networks in spines^26^ that show distances below about 50 nanometers, judged by comparing network dimensions to the diameter of individual actin filaments, ~ 7 nm. Consequently, this analysis suggests that spine stiffness originates from the cross-linking of actin architecture, which can be due to Arp2/3 complexes^29^.

Importantly, because actin remodeling during LTP follows the NMDAR-dependent Ca^2+^ signaling cascade, crosslink-dependent spine stiffness can be expected to correlate with spine morphological changes and synaptic strength, which would explain the observed correlation with spine size (Fig. 4a) and the lognormal-like distribution of spine stiffness (Fig. 4c), as synaptic strength is known to follow a lognormal-like distribution^30^.

Taken together, the extreme stiffness of synapses, attributable to the extensive crosslinking of the postsynaptic F-actin network, along with the biophysical model summarized in Fig. 5, points to an indispensable role stiffness could be playing in synapses: a physical messenger that conveys information about LTP-driven biomolecular and structural changes in the postsynaptic spine to the synaptic adhesion molecules, resulting in stronger synaptic adhesion. This potential coupling of information from the physical domain (stiffness) back to biochemical interactions (adhesion molecules) could link structural plasticity to functional plasticity^31^, and help maintain synaptic connectivity.

## Methods

### Hippocampal neuron culture preparation

Animal work was approved by the Columbia University Institutional Animal Care and Use Committee. Hippocampal neuron cultures were prepared following a modified version of the previously described Brewer method^32^. Fetuses at embryonic day 18 (E18) from timed pregnant Sprague-Dawley rats (Taconic Farms; Hudson, NY) were sacrificed and the hippocampi removed and collected in room temperature Hank’s balanced salt solution (HBSS-; Thermo Fisher Scientific 14025076), supplemented with 0.6% (w/v) glucose (HBSS+). The hippocampi were then incubated in 0.05% trypsin (Thermo Fisher Scientific 25300054) for 15 minutes at 37°C and washed with HBSS+ three times for 10 minutes each. Finally, the neurons were dissociated in Neurobasal medium (Thermo Fisher Scientific 21103-049) supplemented with B27 supplements (Thermo Fisher Scientific 17504-044) and 0.5 mM L-Glutamine (Thermo Fisher Scientific 25030). Neurons were plated at a density of 100,000 cell/mL in glass bottom dishes coated with 1 mg/mL poly-L-lysine (Sigma-Aldrich P2636) and 10 μg/mL mouse protein laminin (Thermo Fisher Scientific 23017-015). 50 mm glass bottom dishes (WillCo GWSt-5040) were used in most experiments unless otherwise specified. The resulting neuronal cultures consisted of a population enriched in large pyramidal neurons. Cultures were maintained in 5% CO_2_ humidified incubator at 37°C and used after 14-25 days *in vitro* (DIV) to image mature synapses, and DIV 5-7 to image immature protrusions. Before imaging, culture medium was replaced with Tyrode’s buffer (125 mM NaCl, 2 mM KCl, 3 mM CaCl_2_, 1 mM MgCl_2_-6H_2_O, 10 mM HEPES, 30 mM D-glucose, adjusted to 300 mOsm with sucrose, pH adjusted to 7.4 with NaOH)^33^ at room temperature. Neuronal activities were verified with calcium indicator Oregon Green (Thermo Fisher Scientific O6807) showing neuron firing.

### Torsional harmonic cantilevers

T-shaped cantilevers were custom made^8^ by Bruker-Nano, Inc with the following specifications: the cantilever bodies were made of silicon nitride and the tip was made of silicon. The length, width, and thickness of the cantilevers were nominally 85 μm, 9 μm, and 650 nm. The width at the free end was 60 μm, and the tip offset 25 μm, tip height 5 μm or 6.5 μm. Cantilevers were coated with silicon nitride via plasma-enhanced chemical vapor deposition to a radius of 75nm or 100 nm. Flexural and torsional deflection sensitivities of cantilevers were determined from ramp plots, assuming flexural and torsional motions to be described by springs in series. The spring constants of flexural (approximately 0.2 N/m) and torsional (approximately 1.0 N/m) deflections were determined from the respective thermal noise spectra.

### Atomic force microscopy imaging

Glass bottom dishes were mounted on the stage of an inverted fluorescence microscope Axio Observer Z1 (Zeiss) and neurons were perfused with Tyrode’s buffer during imaging. TH-AFM experiments were performed with BioScope Catalyst (Bruker) and imaging was carried out in fluid tapping mode. T-shaped cantilevers were analyzed in real time to create topographical and mechanical maps as previously described^9^. The set point amplitude was approximately 45 nm and the tip-sample interaction force was approximately 300 pN. Elastic modulus was calculated by fitting the force-distance curves with a Derjaguin-Muller-Toporov (DMT) model^34^ with a hemispherical indenter (see Supplementary Methods for additional discussion about elastic modulus calculation). All AFM images were recorded with 512 pixel × 256 pixel, 256 pixel × 256 pixel, or 256 pixel × 128 pixel over areas of 5 μm × 5 μm to 25 μm × 25 μm with a scan rate 0.5 to 1.5 Hz.

### Atomic force microscopy data analysis

AFM data were processed and analyzed with NanoScope Analysis 1.70 (Bruker), SPIP 5.0 (Image Metrology), Gwyddion, and ImageJ. For quantitative measurement, in order to reduce noise, a 3×3 median filter was applied to stiffness images. Due to edge effect, AFM measurement is more reliable on top of a structure close to the center, and less so close to the edge. Thus, we selected areas of interest (AOI) of at least 10 pixel × 10 pixel close to the center on a spine structure identified by the topographical image, and used the maximum value in the AOI to represent the elastic modulus of a spine. We noticed that the stiffness of a dendritic shaft and an immature protrusion did not vary substantially along its length. We thus drew a section line of at least 10 pixels along the length of a shaft structure or an immature protrusion close to its centerline, and used the average value along the section to represent the elastic modulus of a shaft and a filopodium, respectively. For apparent spine size measurement, we applied a median filter to the AFM stiffness image. Substrate signal was measured by selecting a substrate region in the image and plotting a Gaussian distribution histogram showing mean *μ* and standard deviation σ. The threshold was then set as *μ + 3σ* to identify AOI for area measurement. For intensity profile in Fig. 2c, raw elastic modulus data were used. For visualization purpose only, a low pass filter was applied to height images; spike removal with vertical interpolation and local mean equalization were applied to stiffness images.

### Epifluorescence microscopy

Optical images were taken before and after TH-AFM imaging using an inverted microscope (Axio Observer Z1; Zeiss) at different magnifications (10X, 20X, 100X EC Plan-Neofluar, Zeiss). We used phase contrast at 10X and 20X and brightfield at 100X in optical imaging. After live imaging, the location of neurons of interest was marked by labeling the relative position of the perfusor (Bruker) to the dish, and the relative position of perfusor to the objective to ensure the same regions could be captured after immunocytochemistry. Fluorescence images were taken using the same epifluorescence microscope (Axio Observer Z1; Zeiss) with proper filter sets (Zeiss and Chrome) at 100X magnification (1.3 NA). All images were captured with a standard CCD camera (Hamamatsu) at 1344 pixel × 1024 pixel resolution.

### Functional labeling of presynaptic boutons with FM 4–64

At the end of the TH-AFM experiment, neurons were incubated in high KCl Tyrode’s buffer (77 mM NaCl, KCl 50 mM, 3 mM CaCl2, 1 mM MgCl_2_-6H_2_O, 10 mM HEPES, 30 mM D-glucose, adjusted to 300 mOsm with sucrose, pH adjusted to 7.4 with NaOH) with 10 μM FM 4-64 for 45 seconds^35^, and then washed with calcium-free Tyrode’s buffer (125 mM NaCl, 2 Mm KCl, 1 mM MgCl_2_-6H_2_O, 10 Mm HEPES, 30 mM D-glucose, adjusted to 300 mOsm with sucrose, pH adjusted to 7.4 with NaOH) for 15 minutes to remove non-specific membrane bound FM 4-64. After FM 4-64 imaging, cells were washed with normal Tyrode’s buffer for 30 minutes to remove trapped FM dyes, before being proceeded to immunocytochemistry.

### Immunocytochemistry

The primary antibodies used were PSD-95 (1:1,000, mouse; Abcam ab99009) and Synapsin-1 (1:1,000, rabbit; Cell Signaling 5297). The secondary antibodies used were Alexa Fluor^®^ 488 Goat Anti-Mouse (1:5000, Thermo Fisher Scientific A-11029), Alexa Fluor^®^ 488 Goat Anti-Rabbit (1:5000, Thermo Fisher Scientific A-11034), Alexa Fluor^®^ 546 Goat Anti-Mouse(1:5000, Thermo Fisher Scientific A-11030), Alexa Fluor^®^ 546 Goat Anti-Rabbit (1:5000, Thermo Fisher Scientific A-11035), Cy5^®^ Goat Anti-Mouse (1:5000, Thermo Fisher Scientific A10524), Cy5^®^ Goat Anti-Mouse (1:5000, Thermo Fisher Scientific A10523). To label F-actin, Alexa Fluor^®^ 546 Phalloidin (1:500, Thermo Fisher Scientific A22283) was added to the secondary antibody solution. After TH-AFM imaging, neurons were fixed with 4% (w/v) paraformaldehyde (Thermo Fisher Scientific 28908), permeabilized with 0.3% (v/v) Triton X-100 (Sigma-Aldrich 93443) in phosphate-buffered saline (PBS), and incubated with primary antibodies diluted in SuperBlock Blocking Buffer (Thermo Fisher Scientific 37515) overnight at 4°C. Neurons were then incubated with secondary antibodies at room temperature for 30 minutes to 2 hours.

### Sample preparation for TEM

To acquire the ultrastructure of synapses, neurons were cultured in homemade glass bottom dishes (Corning^®^ 60mm TC-Treated Culture Dish 430166) on gridded coverslips (Electron Microscopy Sciences 72264-18 and 72265-50). After live optical and TH-AFM imaging, neurons were fixed in the dish with 2.5% (w/v) glutaraldehyde in 0.15 M sodium cacodylate buffer (pH 7.4) at room temperature for 1 hour and then at 4°C overnight. Neurons were then rinsed 3 times in 0.1 M cacodylate buffer at 4°C and post-fixed with 1% OsO_4_ in 0.1 M cacodylate buffer at 4°C for 1 hour. After block staining with 1% uranyl acetate at 4°C for 1 hour, neurons were rinsed 3 times with ddH_2_O at 4°C and dehydrated in a gradient of ethanol: 30%, 50%, 70% at 4°C for 5 minutes each, 85%, 95% at room temperature for 5 minutes each, and 100% four times for 5 minutes each. Neurons were infiltrated in 100% ethanol/Araldite 502 (Electron Microscopy Sciences 13900) at room temperature: 1:1 twice for 10 minutes each, 1:2 for 10 minutes, and 100% Araldite three times for 10 minutes each, then 100% Araldite overnight. The sample was flat embedded and polymerized at 60°C for 48 hours.

### Serial section TEM

The sample block was detached from the coverslip by immersing the whole dish in liquid nitrogen, and then trimmed under stereoscope. The grid pattern imprinted in the resin served as landmarks. 70 nm serial ultrathin sections were cut using Leica UC6 ultramicrotome (Leica Microsystems Inc., Buffalo Grove, IL), collected on formvar coated slot grids, and stained with uranyl acetate and lead citrate. The top few sections containing the marker grid pattern were recognized under transmission electron microscope (Philips CM-12, FEI, Eindhoven, The Netherland) at lower magnification (170X), and were used to locate the regions of interest based on the comparison of neurites morphology from optical images. To identify regions of interest in deeper sections, the relative location of regions of interest on TEM sections was marked on the captured images (Gatan 4k×2.7k digital camera, Gatan Inc., Pleasanton, CA) and used as reference. Serial sections of neurites and synapses were then imaged at 170X - 66000X magnification.

### Image processing

Optical images were processed and quantified with ImageJ and Caltracer 2 (available through: http://blogs.cuit.columbia.edu/rmy5/methods/). Background signal was measured by selecting dark regions in an image and plotting a Gaussian distribution histogram showing mean *μ* and standard deviation *σ*. The intensity of the whole image was then subtracted by *μ + 3σ* to remove noise. Subtraction processed images were then analyzed using Caltracer to identify colocalization. Puncta contours of each marker were detected using an automated algorithm based on fluorescence intensity, puncta size, and shape, and were adjusted by visual inspection. Contours of different markers (PSD-95 and Synapsin-1) were overlaid and a threshold of 5% overlap was used for all potential colocalization detection. Colocalized contours were counted and overlaid with stiffness images. TEM images were processed with the FIJI plugin Enhance Local Contrast (CLAHE) (available through: http://imagej.net/Enhance_Local_Contrast_(CLAHE)) to enhance local contrast for visualization, with histogram bins 50. Optical images, AFM images, and TEM images of different magnifications of the same area were aligned in Adobe Photoshop and visually inspected. In particular, due to the non-linear lens distortions induced by the electromagnetic lenses of TEM, serial section TEM images were usually distorted. In order to align TEM images with optical images and AFM images, we skewed the TEM images with shear transformation in Adobe Photoshop. For visualization purpose only, brightness and contrast was adjusted, median filter was applied, and pixel number was increased to smoothen the pixelated images in Adobe Photoshop. For quantitative analysis including intensity profiles, raw data were used unless otherwise specified.

### Statistical analysis

Error bars in all figures represented standard error of mean. Two-tailed tests were used in statistical analysis. Sample size (n), p value, and Pearson correlation coefficient (r) were given in figure legends when applicable. Each neuron culture represented an independent experiment. A significance level of 0.01 was used in hypothesis tests and p < 0.01 was considered significant. To fit the transformed data to a normal distribution, the maximum likelihood estimation method was used. Goodness of fit tests of the transformed data in a normal distribution were performed using two-tailed Kolmogorov-Smirnov tests with the null hypothesis that the transformed data is normally distributed. We tested goodness of fit for different transformations including logarithm, square root, and cubic root, as well as for non-transformed data (see Supplementary Table 5 for p values). Given that for spine stiffness (kPa), shaft stiffness (kPa), spine size (μm^2^), logarithm transformation has the highest p value (the probability of the null hypothesis being true), the data is likely to follow a lognormal-like distribution. All data were analyzed using Python.

## Data availability

The data supporting the findings of this study are available within the paper and the Supplementary Information. The remaining data are available from the corresponding author on request.

## Acknowledgements

We thank NYU Microscopy Core, Kristen Dancel-Manning, Chris Petzold, and Alice Liang, for their assistance in TEM, and Reka Recinos and Hardik Patel for neuronal cultures. This work was supported by the NIH Director’s New Innovator Award Program (1DP2-EB018657), the NIMH (R01MH101218, R01MH100561), NINDS (R01NS110422; R34NS116740), and the U.S. Army Research Laboratory and the U. S. Army Research Office under contract number W911NF-12-1-0594 (MURI).

## Author Contributions

J.Y. carried out AFM imaging, fluorescence microscopy imaging, and TEM imaging, contributed to the conceptualization, cantilever preparation, improvement of imaging protocols, data analysis, and development of mechanical models. N.M. demonstrated proof of principle for the AFM imaging in neurons and contributed to the development of imaging protocols. S.G.H. contributed to the development of mechanical models and data analysis. J.J.M. contributed to AFM and fluorescence microscopy imaging, and cantilever preparation. R.Y. contributed to the conceptualization, experimental design, and discussions. R.L. prepared neuron cultures, contributed to the conceptualization, experimental design, and discussions. O.S. contributed to the conceptualization, supervision, cantilever design, development of mechanical models, and data analysis. J.Y. and O.S. wrote the manuscript with input from all coauthors.

