## Supplementary Information for "Extreme stiffness of neuronal synapses and implications for synaptic adhesion and plasticity"

### Supplementary Notes

**Contact mechanics model in stiffness calculation**

Elastic modulus was calculated by fitting the AFM force-distance curves using a Derjaguin-Muller-Toporov (DMT) model[1] with a hemispherical indenter as previous described[2]. The interaction force during AFM indentation is written as the following:

$$F = F_{adh} + \frac{4}{3}E^{*}\sqrt{R^{*}}d^{\frac{3}{2}} \quad (1)$$

$F$ denotes the tip-sample interaction force. $F_{adh}$ denotes a constant adhesion force measured by the peak negative force during AFM retraction. $E^{*}$ denotes the effective elastic modulus. $d$ denotes the indentation depth. $R^{*}$ denotes the effective radius:

$$\frac{1}{R^{*}} = \frac{1}{R_{tip}} + \frac{1}{R_{sample}} \quad (2)$$

$R_{tip}$ is the tip radius, and $R_{sample}$ is the sample radius. Because $R_{sample}$ (spine and shaft radius) is relatively large compared to $R_{tip}$, we neglected $R_{sample}$ in the stiffness calculation in Fig. 4a for 409 spines and shafts.

**Sensitivity of spine stiffness - spine size correlation to contact mechanics models**

Our results in Fig. 4c showed spine stiffness was correlated with spine size. It is important to consider whether such correlation is introduced because the calculation of $E^{*}$ depends on spine radius $R_{sample}$ (See equations (1,2)). In order to analyze the correlation between spine stiffness $E^{*}$and spine size $R_{sample}$, we first took into consideration $R_{sample}$ to calculate $E^{*}$in the DMT model (See equations (1,2)). To estimate $R_{sample}$, we first measured the apparent spine size from AFM stiffness images by thresholding the stiffness signal to identify region of interest for area measurement. We used area, $S$, to represent the apparent spine size in Fig. 4b, c. Assuming the measured area is a round flat surface, we estimated sample radius from $S = \pi {R_{sample}}^2$. We used the DMT model with $R_{sample}$ in the stiffness calculation in Fig. 4c. Inclusion of $R_{sample}$ in the DMT model slightly increases measured spine stiffness, suggesting that our stiffness calculation in Fig. 4a may be an underestimate of sample stiffness.

When the AFM tip exerts force onto the spine head, the forces are transmitted to the spine head – substrate interface. Deformations of the spine head at this interface during interactions with AFM tip could also affect stiffness measurements. Because spine diameters are large compared to the AFM tip, neglecting the spine-substrate interface in the DMT model is a plausible assumption. Furthermore, presence of adhesive forces between the substrate and the spine head would increase the apparent stiffness of this interface. However, it is still possible to make a worst-case estimate of the contributions from the spine-substrate interface by assuming that there are no adhesive forces at this interface and the spine head is making a sphere-plane contact with the substrate.

We carried out this worst-case analysis by considering indentations on both surfaces: $d = d_{top} + d_{bottom}$. $d_{top}$ is the indentation depth on the top surface of the sample, and $d_{bottom}$ is the indentation depth on the bottom of the sample close to the substrate. In the DMT model used in Fig. 4a, we assumed the indentation between the bottom of the sample (spine head) and the substrate is trivial and thus used $d = d_{top}$. Here, to account for both $d_{top}$ and $d_{bottom}$ to make an worst-case estimate of elastic modulus considering sample geometry, we adapted the "sphere between two parallel planes" model[3]. Given $F$ is the same on both surfaces, we could write $F = \frac{4}{3}E^*\sqrt{R^*_{top}}d_{top}{}^{\frac{3}{2}}$, and $F = \frac{4}{3}E^*\sqrt{R^*_{bottom}}\,d_{bottom}{}^{\frac{3}{2}}$, thus

$$\frac{d_{bottom}}{d_{top}} = \sqrt[3]{\frac{R^*_{top}}{R^*_{bottom}}} \tag{3}$$

From equation (2), we could get $R^*_{bottom} = R_{sample}$, and $R^*_{top} = \frac{R_{tip} \times R_{sample}}{R_{tip} + R_{sample}}$. Thus equation (3) can be written as:

$$\frac{d_{bottom}}{d_{top}} = \frac{1}{\sqrt[3]{1 + \frac{R_{sample}}{R_{tip}}}} \tag{4}$$

From equation (1), the worst-case elastic modulus $E^*_2$ can be written as $F = \frac{4}{3}E^*_2\sqrt{R^*_{top}}d_{top}{}^{\frac{3}{2}}$ while the DMT model $E^*$ can be written as $F = \frac{4}{3}E^*\sqrt{R^*_{top}}(d_{top} + d_{bottom})^{\frac{3}{2}}$. We could then derive $E^*_2$ as following:

$$E^*_2 = E^* \times \left(1 + \frac{1}{\sqrt[3]{1 + \frac{R_{sample}}{R_{tip}}}}\right)^{\frac{3}{2}} \tag{5}$$

Using this worst-case $E^*_2$ which accounts for sample geometry $R_{sample}$ and sample-substrate interaction, we performed correlation analysis, and revealed that the worst-case $E^*_2$ is still correlated with spine size with r = 0.2595 and p = 2.461E-03. In the DMT model with $R_{sample}$ (spine radius) taken into account, $E^*$ is correlated with spine size with r = 0.3837 and p =

4.719E-06 (model in Fig. 4b, c). In the DMT model without $R_{sample}$ (spine radius) taken into account (i.e. assuming $R_{sample} \gg R_{tip}$), $E^*$ is correlated with spine size with r = 0.4384 and p = 1.174E-07 (model in Fig. 4a). In all three models, spine stiffness is correlated with spine size.

**Contact mechanics model of elastic modulus and synaptic adhesion**

To make an order of magnitude estimate of the minimal elastic modulus of a spine required to maintain morphology in the presence of adhesive force, we used a contact mechanics model to relate deformation of contacting structures to adhesive force and elastic modulus. We treated the spine-bouton system as a pliable ball (spine head) pressing against a relatively flat and hard surface (bouton). Assuming that the spine head is deformed by the adhesive force at synapse and that no significant adhesive interaction occurs outside of the active zone, we used the Johnson-Kendall-Roberts (JKR) model[4] to estimate the minimal elastic modulus required. We assumed that the shape of the spine is deformed significantly when the diameter of the contact zone becomes comparable to the spine radius. Therefore, we determined the required elastic modulus to prevent the contact diameter from becoming larger than spine radius. According to the JKR model, contact radius (half the diameter) $a$ could be written as the following:

$$a^3 = \frac{3R}{4E^*}\left(P + 3\gamma\pi R + \sqrt{6\gamma\pi RP + (3\gamma\pi R)^2}\right) \quad (6)$$

Here, $R$ denotes the radius of curvature of a typical spine head, $E^*$ denotes the effective elastic modulus, $P$ denotes the applied load, and $\gamma$ denotes the work of adhesion. Note that there could be pulling force across a synapse[5], thus $P$ is likely to be negative. However, the adhesive force must be significantly larger than the applied load $P$ to hold the pre- and postsynapses together. Therefore, we neglected $P$ in our model. We further assumed that the effective elastic modulus

$E^*$ primarily comes from the stiffness of the spine. This is because axon is under tension[5], which could help maintain bouton's shape. Furthermore, the adhesion molecules on the presynaptic side could be ultimately connected to microtubules, which are resistant to deformation. We thus neglected bouton's elastic modulus, and derived the elastic modulus of the dendritic spine $E^*$:

$$E^* = \frac{9\pi R^2}{2a^3}\gamma \tag{7}$$

We assumed that the maximal contact radius at synapse interface to maintain the structural integrity of the spine is half of the spine radius, i.e. $a = \frac{R}{2}$, and obtained the relationship between the minimal elastic modulus of the spine head $E^*$ and the surface energy $\gamma$:

$$E^* = \frac{36\pi}{R}\gamma \tag{8}$$

To determine the minimal elastic modulus, we need to estimate the work of adhesion $\gamma$, which in our model corresponds to the adhesion energy between pre- and postsynapses. For this, we considered the adhesion mediated by synaptic adhesion molecules. We modified the model developed by Chen et al.[6], which characterizes the adhesion mediated by pairs of adhesion molecules. According to Chen et al., the adhesive energy $\Delta G$ depends on the number of adhesion molecule dimers formed between two cells and the free energy of the monomer-dimer reaction. Assuming a local chemical equilibrium at cell-cell interface, the free energy can be calculated from dissociation constant $K_d$, which is given by the concentrations of monomers and dimers. The 3D concentration of adhesion molecules could be converted from the 2D surface density using the "interfacial shell" model purposed by Chen et al.. Taken together, adhesive energy $\Delta G$ at synapse can be written as the following:

$$\Delta G = AhN_A \frac{C^2}{K_d} RTln(K_d) \tag{9}$$

Here, $A$ denotes total surface area where the adhesion molecules reside, i.e. active zone, $h$ denotes the shell thickness in the "interfacial shell" model, and we used the thickness calculated by Chen et al., 12 nm. $N_A$ is the Avogadro number. $C$ represents the concentration of monomers at synapse. $\Delta G$ has a unit of J/mole. We converted $\Delta G$ to surface energy $\gamma$ in the unit J/m$^2$ by considering the number of molecules and the surface area at the active zone of the synapse:

$$\gamma = \frac{N}{N_A} \times \frac{\Delta G}{A}. \tag{10}$$

Here, $N$ denotes total number of one type of adhesion molecules on the membrane. In our model, we considered the following 2 types of adhesion molecules: N-cadherin and NCAM-140, because they are widely-studied synaptic adhesion molecules and their dissociation constants have been measured. We estimated the number of each molecule at synapse based on mass spectrometry data[7] and used the morphological characteristics of an average synapse from electron-microscopy 3D reconstruction data[8,9].

We calculated the surface energy to be 3.43E-04 J/m$^2$. From equation (8), we obtained the minimal effective elastic modulus of the spine head, 183 kPa. See Supplementary Table 1 - 4 for values used in this model.

### Supplementary Figures

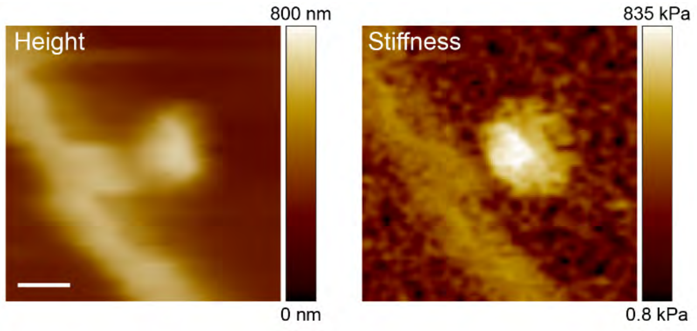

**Supplementary Figure 1: TH-AFM images of a stiff synapse-like structure.**

The AFM height and stiffness images of the synapse-like structure in Fig. 1c. The elastic modulus values of the stiff structure and the shaft are 509.0 kPa and 42.9 kPa, respectively. Scale bar: 1 µm.

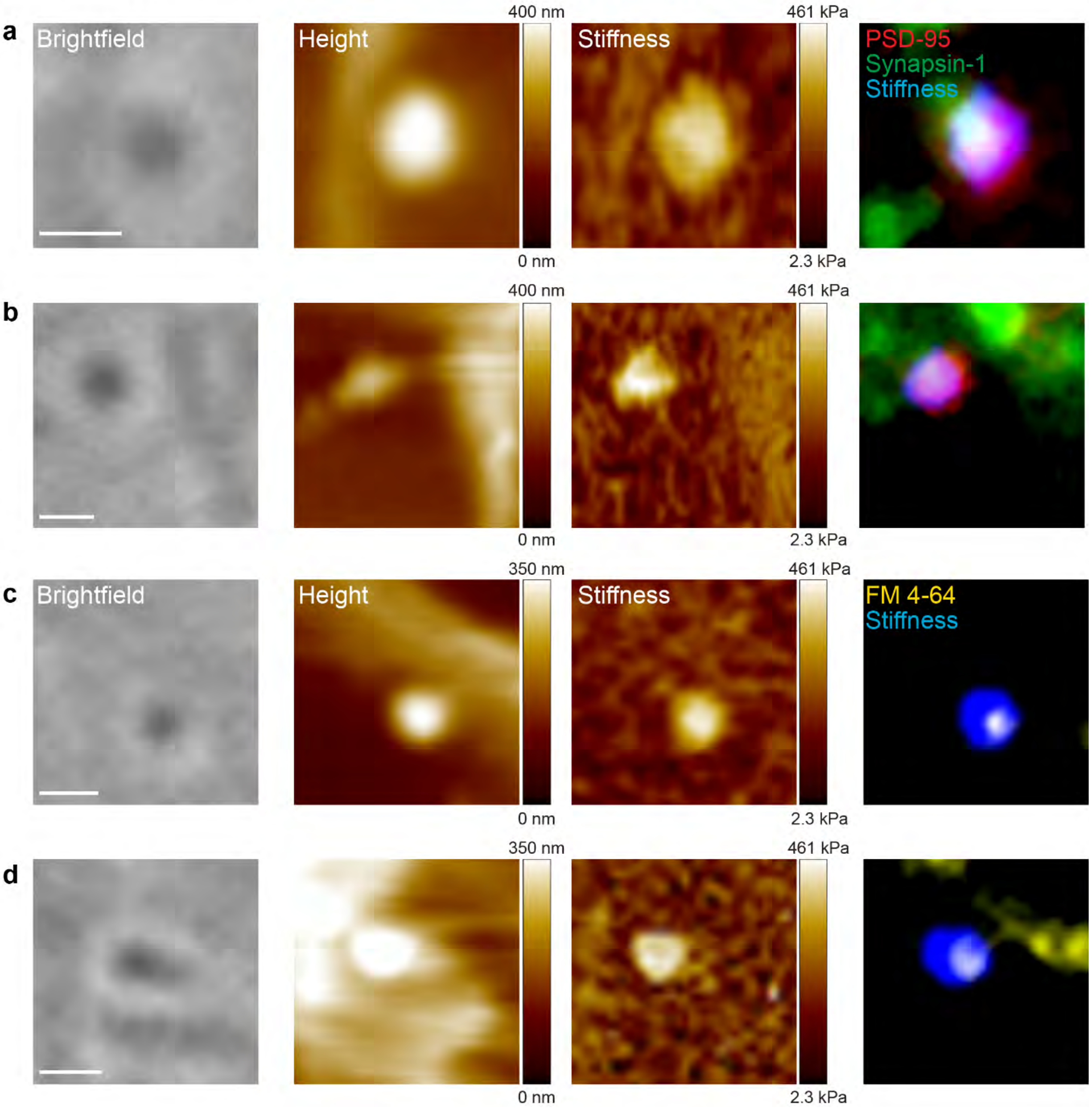

**Supplementary Figure 2: Stiff synapse-like structures are labeled with synaptic markers.**

**a, b** Aligned brightfield, AFM height, AFM stiffness, and immunofluorescence images of two representative stiff synapses labeled with both synaptic markers. 263 synapses from 20 neuron cultures were imaged with TH-AFM and aligned with immunofluorescence images. Threshold was applied to the stiffness image colored in blue. The elastic modulus values of 2 synapses are 271.3 kPa and 363.5 kPa, and the elastic modulus values of shafts are 37.3 kPa and 30.9 kPa. **c, d**

Aligned brightfield, AFM height, AFM stiffness, and fluorescence images of two representative stiff synapses labeled with FM 4-64. 97 synapses from 7 neuron cultures were imaged with TH-AFM and aligned with FM images. The elastic modulus values of 2 synapses are 310.8 kPa and 190.7 kPa, and the elastic modulus values of shafts are 24.6 kPa and 26.6 kPa. Scale bar: 500 nm.

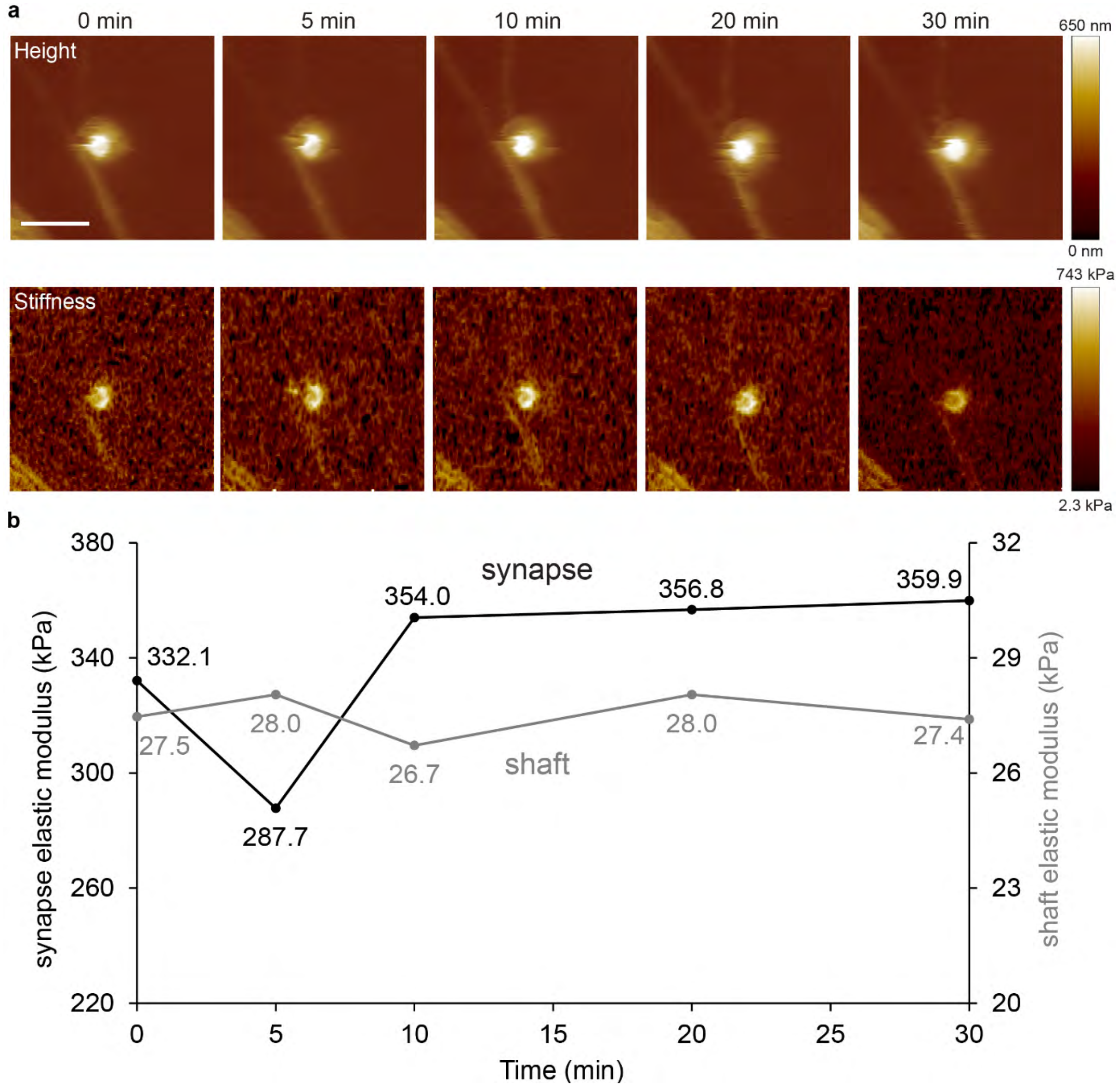

**Supplementary Figure 3: Synapse stiffness does not vary significantly during imaging.** **a** Time-lapse TH-AFM height and stiffness images of a synapse. The same area was scanned with TH-AFM at 0, 5, 10, 20, 30 min. Scale bar: 2 μm. **b** Stiffness of the synapse (black) and shaft (grey) did not change drastically over 30 minutes. The stiffness of the synapse dropped to 287.7 kPa from 332.1 kPa at 5 min (13.4% decrease compared to 0 min) and increased to 354.0 kPa at 10 min (6.6% increase compared to 0 min). These variations were small and could probably be due to measurement uncertainty.

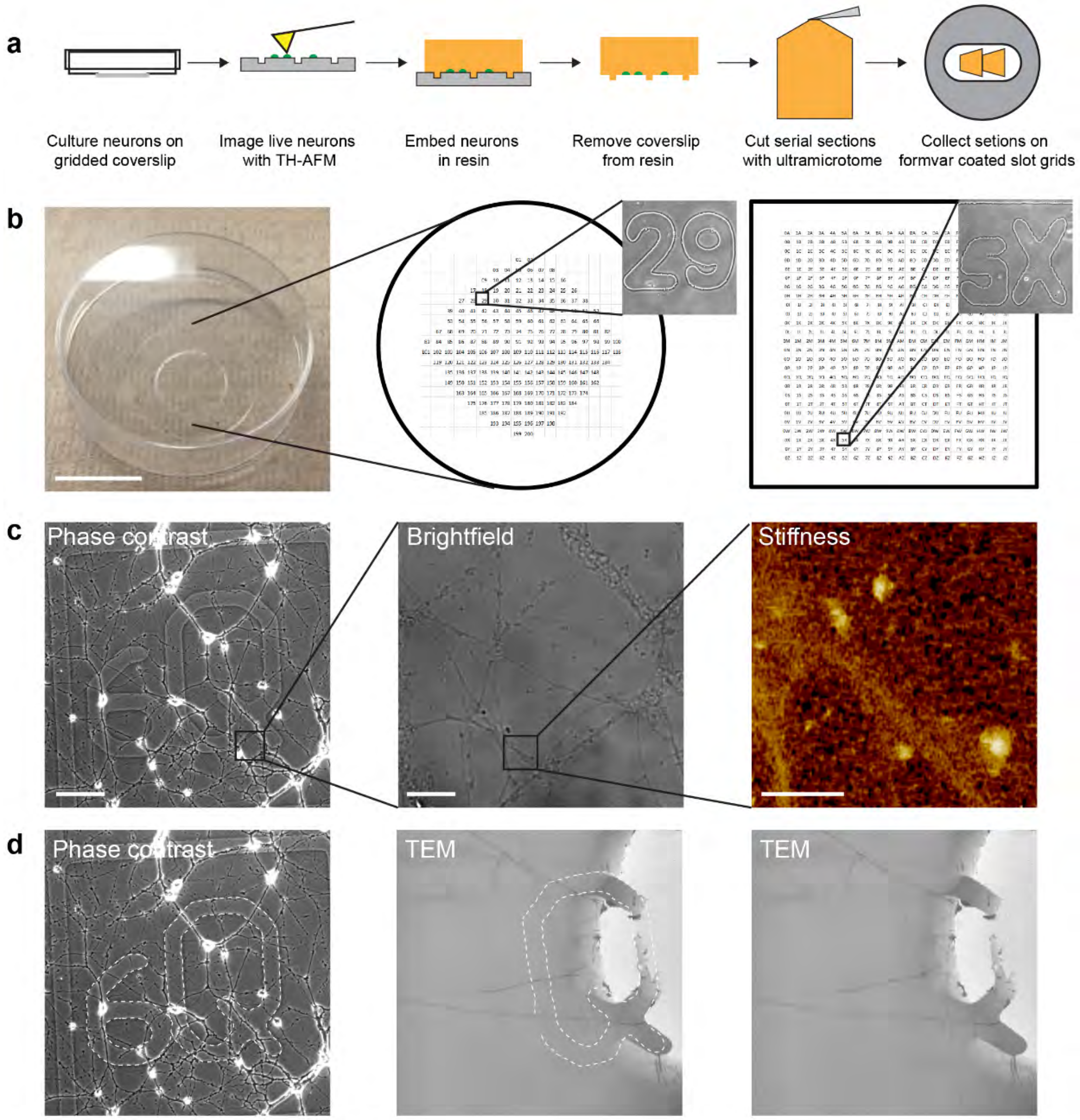

**Supplementary Figure 4: Correlative TH-AFM/TEM imaging.**

**a** To correlate AFM, TEM, and optical images of the same synapses, neurons (green) were cultured in homemade glass bottom dishes with gridded coverslips. After TH-AFM imaging, neurons were fixed, stained, and embedded in resin (orange). The sample blocks were detached from the coverslip, trimmed to 70 nm serial ultrathin sections, and collected on formvar coated slot grids. **b** A 60 mm petri dish with a gridded coverslip attached to the bottom used for neuron cultures. Schematic images of two types of gridded coverslips used in the experiment: numeric and alphanumeric pattern. Phase contrast images of pattern “29” and “5X" are shown here as examples. Scale bar: 2 cm. **c** Alphanumeric pattern “6Q” was recognized under optical microscope in neuron cultures. AFM stiffness image of the boxed area in the high magnification optical image is shown. Scale bar: phase contrast 100 μm, brightfield 10 μm, AFM stiffness 2 μm. **d** The grid pattern observed in the optical phase contrast image and later imprinted in the resin served as landmarks. Marked dashed lines show the pattern “6Q”. The marker grid pattern was recognized in the top TEM section at lower magnification, and was used to locate the regions of interest based on the comparison of neurites morphology from optical images. Note that only part of “Q” was visible in the TEM image possibly due to the cutting angle in serial section TEM sample preparation.

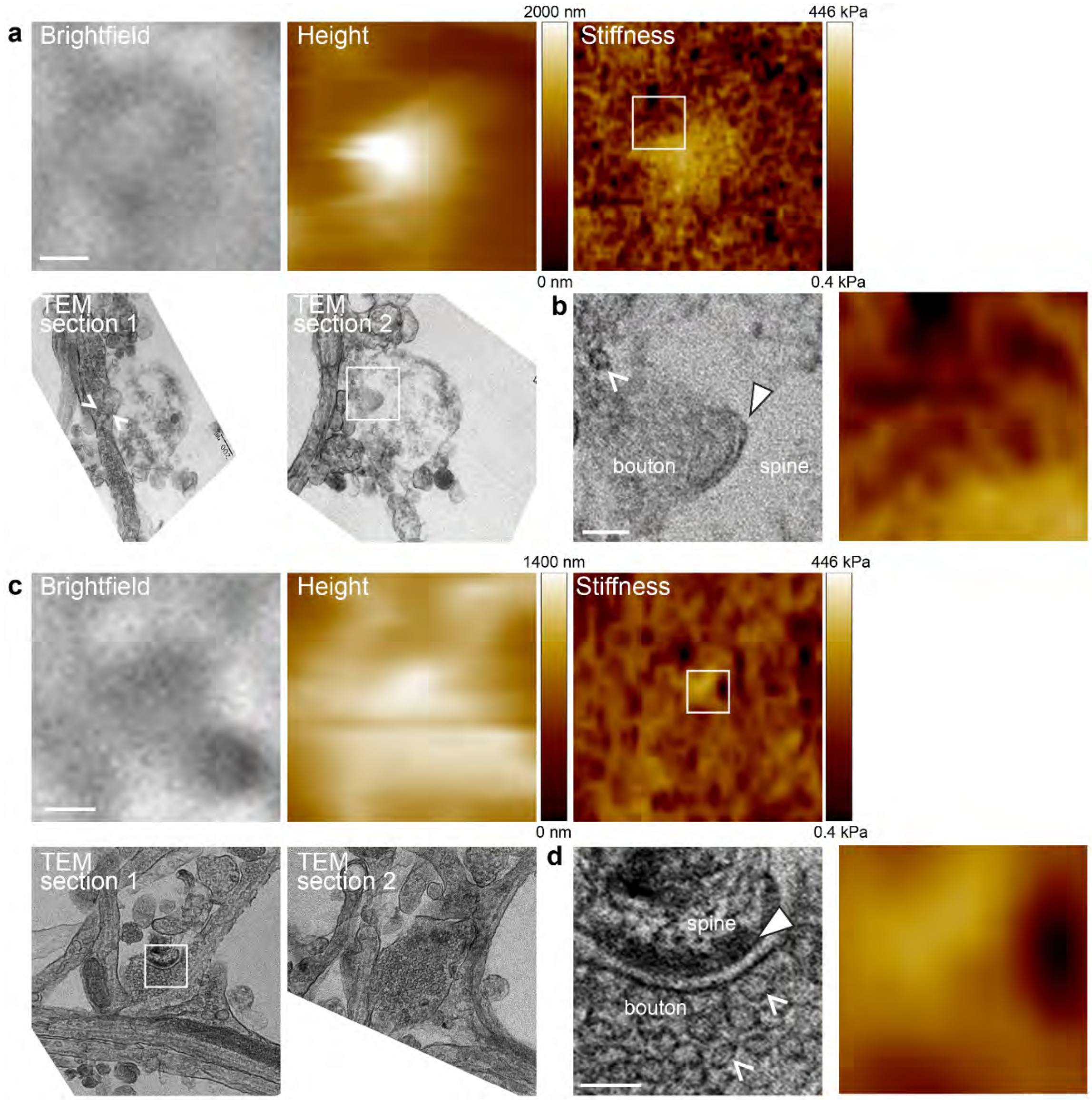

**Supplementary Figure 5: Examples of correlative TH-AFM/TEM images of synapses.**

**a** Aligned brightfield, AFM height, AFM stiffness, and serial section TEM images of the same synapse. The bouton containing vesicles appeared in TEM section 1, and the spine head with postsynaptic density appeared in TEM section 2. **b** Zoomed-in TEM image and stiffness image from the boxed areas in the TEM section 2 and AFM stiffness images in **a**. Elastic modulus values of the spine and shaft are 53 kPa and 11 kPa, respectively. Note that high stiffness overlaid with the bottom half of the spine head, while the top half of the spine head did not show topographical features and was likely covered by compliant structures. **c** A bouton containing vesicles formed a synapse with a spine head. Elastic modulus values of the spine and shaft are 26 kPa and 8 kPa, respectively. **d** Zoomed-in TEM image and stiffness image from the boxed areas in the TEM section 1 and AFM stiffness images in **c**. White arrowheads point to the postsynaptic density. White carets point to presynaptic vesicles. Scale bar a, c: 500 nm, b, d: 100 nm.

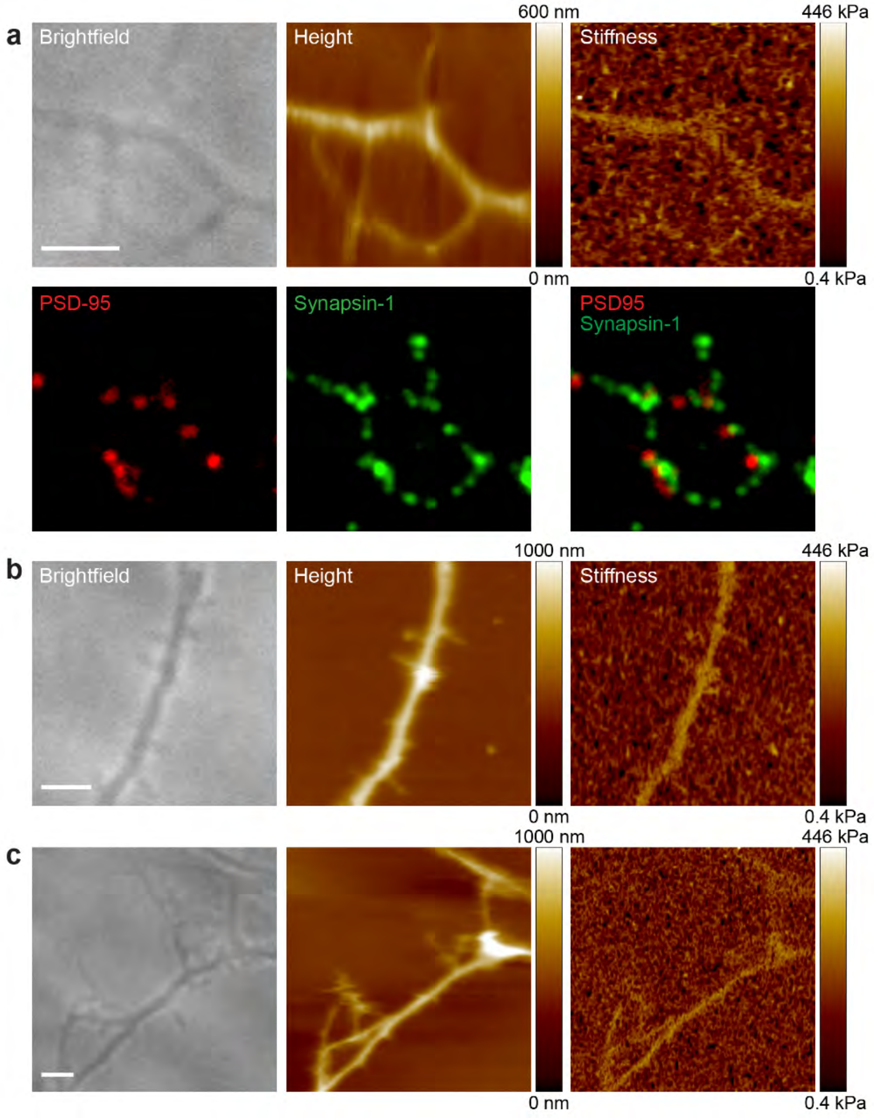

**Supplementary Figure 6: Immature protrusions are not stiff.**

**a** Aligned brightfield, AFM height, AFM stiffness, and immunofluorescence images of neurons on DIV7. Note that the thin protrusions were not highly stiff (14 kPa). **b, c** Aligned brightfield, AFM height, and AFM stiffness images of neurons on DIV6. Elastic modulus values of these non-stiff protrusions are 18 kPa and 14 kPa, respectively. Scale bar: 2 µm. See Methods for quantitative stiffness measurement of areas of interest.

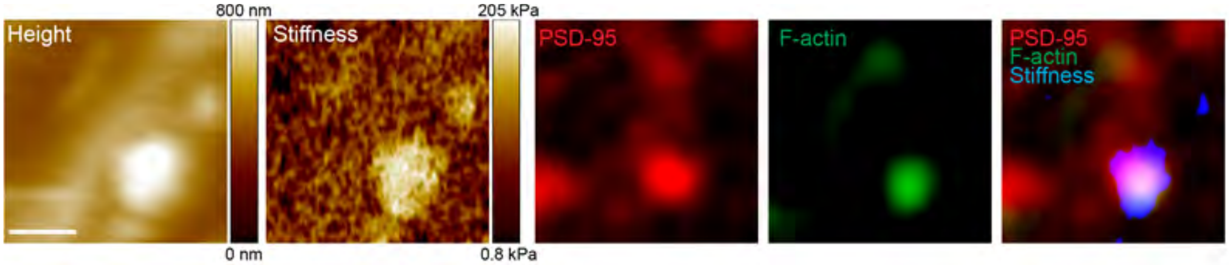

**Supplementary Figure 7: Stiff spines contain enriched F-actin.**

Aligned AFM height, AFM stiffness, and fluorescence images of the same area. F-actin (green) was enriched in the stiff spine (200 kPa), not in the dendritic shaft (19 kPa). Threshold was applied to the stiffness image colored in blue. Scale bar: 500 nm

### Supplementary Tables

| Characteristic | Symbol | Value |
|---|---|---|
| Active zone area | $A$ | 0.07 µm[9] |
| Spine volume | $V' = \frac{4}{3}\pi R^3$ | 0.04 µm[8] |
| Spine radius | $R$ | 0.212 µm |

**Supplementary Table 1: Morphological characteristics of an average synapse.**

| **Molecules at PSD** | **Abundance index by mass spectrometry of PSD[7]** | **Number of molecules per PSD** |
|---|---|---|
| PSD-95 | 24.2 | 300 |
| N-cadherin | 2.5 | 31 |
| NCAM-140 | 1.1 | 14 |

**Supplementary Table 2: Number of adhesion molecules at synapse.**

In our model, we considered only the following 2 types of adhesion molecules: N-cadherin and NCAM-140, because they are widely studied synaptic adhesion molecules and their dissociation constants ($K_d$) have been measured. We estimated the number of each molecule at synapse based on mass spectrometry data from literature.

| | **N-cadherin**[10] | **NCAM-140**[11] |
|---|---|---|
| dissociation constant $K_d$ (M) | 2.58E-05 | 5.5E-05 |
| species | mouse | mouse |
| method | analytical ultracentrifugation | surface plasmon resonance |

**Supplementary Table 3: Dissociation constant of synaptic adhesion molecules.**

| Parameter | Symbol | N-cadherin | NCAM-140 |
|---|---|---|---|
| number per synapse | $N$ | 31 | 14 |
| dissociation constant | $K_d$ (M) | 2.58E-05 | 5.5E-05 |
| free energy | $\Delta g(i,j)$ (J/mole) | -2.62E+04 | -2.43E+04 |
| 2D density on PSD | $\rho$ (/μm$^2$) | 442.74 | 194.81 |
| interfacial shell thickness | $h$ (nm) | 12 | 12 |
| surface area of a single pair of adhesion molecules | $A_c$ (μm$^2$) | 2.26E-03 | 5.13E-03 |
| interfacial shell | $V$ (L) | 2.71E-20 | 6.16E-20 |
| 3D effective concentration | $C$ (μM) | 61.27 | 26.96 |
| monomer concentration | $C_i$ (μM) | 28.90 | 19.82 |
| dimer concentration | $C_{ij}$ (μM) | 32.37 | 7.14 |
| number of monomers | $N_i$ | 14.62 | 10.03 |
| number of dimers | $N_{ij}$ | 16.37 | 3.61 |
| free energy on surface | $\Delta G(I,J)$ (J/mole) | 4.29E+05 | 8.78E+04 |
| free energy on surface | $W$ (J) | 2.21E-17 | 1.99E-18 |
| surface energy at interface | $\gamma$ (J/m$^2$) | 3.15E-04 | 2.84E-05 |

**Supplementary Table 4: Adhesive energy at synapse.**

Total surface energy at synapse interface from these 2 types of adhesion molecules is 3.43E-04 J/m$^2$. Given there are many other types of adhesion molecules at synapse, this theoretical value is likely to be an underestimate of the actual surface energy at synapse.

| | spine stiffness (kPa) | shaft stiffness (kPa) | spine size ($\mu m^2$) |
|---|---|---|---|
| non-transformed | 2.01E-05 | 2.45E-05 | 0.0001 |
| logarithm | 0.3192 | 0.2446 | 0.9486 |
| square root | 0.0231 | 0.0061 | 0.0941 |
| cubic root | 0.1609 | 0.0233 | 0.4158 |

**Supplementary Table 5: p values for Kolmogorov-Smirnov tests of transformed data.**

Null hypothesis: transformed data is normally distributed. Alternative hypothesis: transformed data is not normally distributed. We calculated goodness of fit using two-tailed Kolmogorov-Smirnov tests for different transformations: logarithm, square root, and cubit root. Given that for spine stiffness (kPa), shaft stiffness (kPa), spine size ($\mu m^2$), logarithm transformation has the highest p value (the probability of the null hypothesis being true), data is likely to follow a lognormal-like distribution.
